# Sustained release Resolvin D1 liposomes are effective in the treatment of osteoarthritis in obese mice

**DOI:** 10.1101/2023.01.21.525015

**Authors:** Ameya A. Dravid, Kaamini M. Dhanabalan, Soumyadeep Naskar, Akshi Vashistha, Smriti Agarwal, Bhagyashree Padhan, Mahima Dewani, Rachit Agarwal

**Author notes:** **Corresponding author**, **Email**, **Complete postal address:** 3^rd^ Floor, Biological Sciences Building, Indian Institute of Science, Bangalore-560012, Karnataka, India.

## Abstract

Osteoarthritis (OA) is the most common joint disorder and currently affects > 500 million patients worldwide, with ~60% of them also suffering from obesity. There is no drug approved for human use that changes the course of OA progression. OA is one of the most common comorbidities of obesity, and obesity-related OA (ObOA) is a serious health concern because it shows heightened severity of tissue damage and also predominantly affects the working population. Unresolved inflammation is a major driver of ObOA, thus, resolving disease-associated inflammation is a viable strategy to treat ObOA. Resolvins are highly potent molecules that play a role in the resolution of inflammation and promote tissue healing. However, small molecules (like Resolvin D1; RvD1) have to be administered frequently or prior to injury because they lose their *in vivo* activity rapidly either by lymphatic clearance, or oxidation-mediated deactivation. In this study, we have encapsulated RvD1 in liposomes and established its efficacy in the mouse model of ObOA at much lower dosages than freely administered RvD1. Liposomal RvD1 (lipo-RvD1) acted as a source of the RvD1 molecules for ~11 days *in vitro* in synovial fluid derived from patients. When administered prophylactically or therapeutically, lipo-RvD1 suppressed cartilage damage in male C57BL/6 mice compared to untreated and free RvD1 treatments. This efficacy was achieved by increasing the proportion of the proresolution M2 macrophages over proinflammatory M1 macrophages in the synovial membrane. These results show the potential of lipo-RvD1 as an anti-OA agent.

**Graphical abstract:**
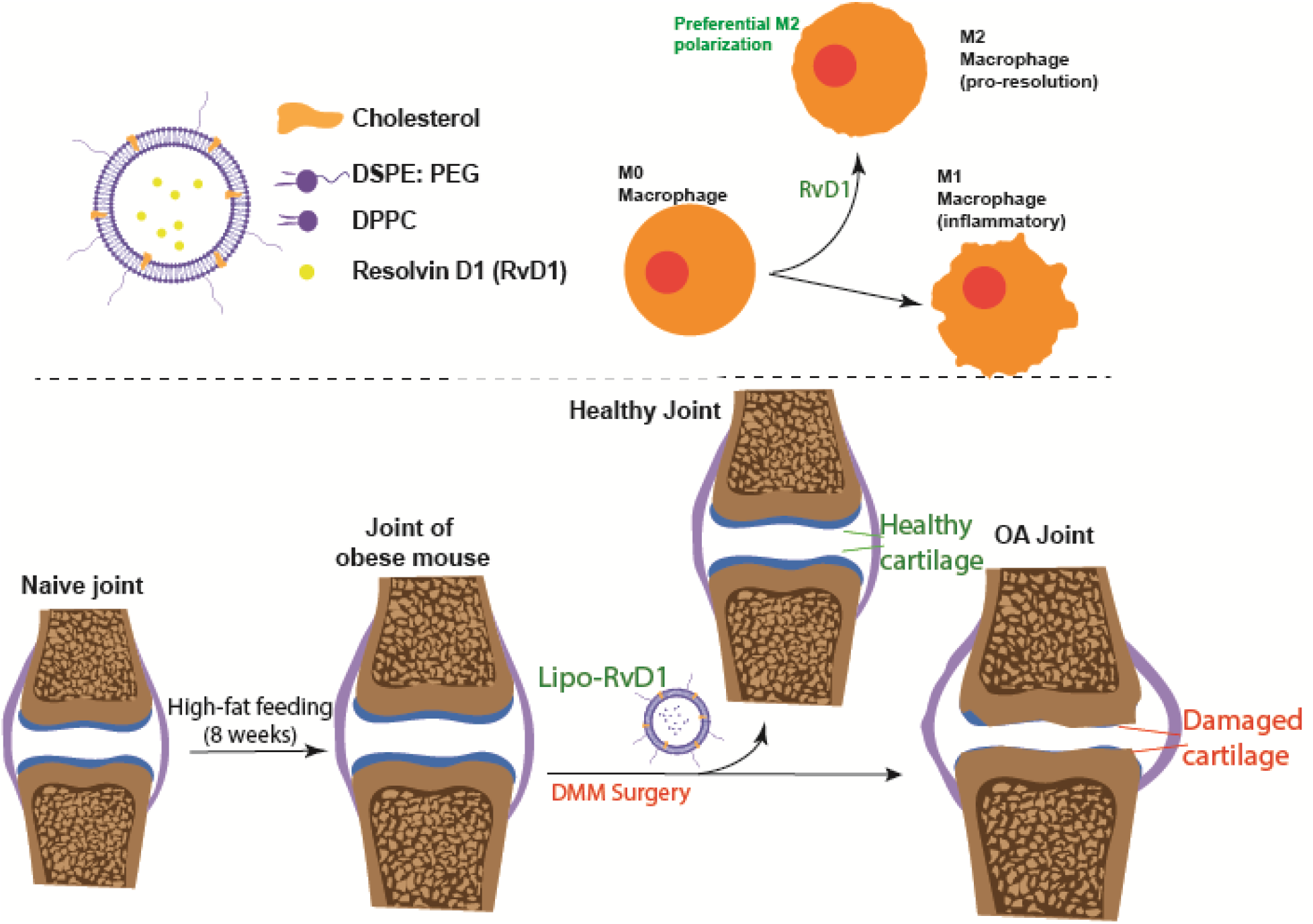
Mechanism of working of lipo-RvD1 in ObOA joint.

## Introduction

Osteoarthritis (OA) is a highly prevalent disease of the articulating joints. Recent reports show that >500 million patients worldwide are affected by OA^1^. OA is known to progressively degrade cartilage and the subchondral bone. Obesity is also heavily linked with OA, where ~59% of all OA patients are also obese^2^. Obesity actively contributes to the progressing cartilage damage via excessive joint loading and elevated inflammatory signaling to subsequently produce an accelerated pathology^3^. Current treatment for OA focuses only on palliative care by administration of analgesics without improving joint function^4^. Steroidal analgesics have several drawbacks like chronic chondrotoxicity and cartilage thinning and are not recommended for long-term administration^5^. Obesity is also characterized by systemic dyslipidemia and the generation of hypertrophic adipocytes and fat depots, which dysregulate the release of several adipokines like resistin, leptin, and visfatin^6^, of which impairment of leptin-mediated signaling is a major contributor to OA^7^. Such impaired adipokine-mediated signaling triggers the release of proinflammatory cytokines like IL-1β and TNF-α in both the systemic circulation and the synovium^7, 8^. Hence, obesity-related metabolic dysregulation contributes to the prevailing chronic OA-associated inflammation and increases tissue damage^6^, and targeting this inflammation is hence a viable strategy to treat ObOA.

Previous attempts that delivered anti-inflammatory molecules to the site of damage have not succeeded in translating into therapy for humans^9, 10^ despite the encouraging results in small animal models^11, 12^, Synovial vasculature rapidly clears small molecules, while the lymphatics drain away macromolecules^13, 14^, resulting in short half-lives in the joint (1-4h) for commonly used steroids^4, 15^. Such molecules, when encapsulated in particles, show increased intraarticular retention because the relatively larger size of the particles prevents lymphatic drainage-mediated clearance^16^. One of the preferred particle-based drug carriers are liposomes. One such liposomal formulation containing palmitoylated dexamethasone, Lipotalon^®^, has been approved in Germany for treating OA-related pain in patients. Liposomal carriers have demonstrated favorable properties like low toxicity, biodegradability, stability, flexible synthesis methods, and ability to incorporate cargo with diverse properties (imaging agents, corticosteroids, MMP inhibitors)^17–19^ and are hence ideal carriers of drugs. These particles have been used for the intraarticular delivery of corticosteroids^20, 21^, and can be used for immunomodulation in OA as well^22^. Liposome-encapsulated molecules show a ~10-fold increase in intraarticular (IA) retention than molecules injected directly^16^. Liposomes have a predictable release of the loaded drugs into the extraliposomal space, thus helping in the process of dosing in humans^16^.

Specialized proresolution mediators (SPMs)^23^ are potent molecules that actively reduce inflammatory factors from the site of damage^23–25^ in a process called resolution of inflammation. Chronic inflammatory diseases, including ObOA, are known to have impaired inflammation resolution pathways^26, 27^. Exogenously supplied SPMs like RvD1 reduce the severity of OA, but strategies involving the direct intraarticular injection of such molecules require high doses and administration of doses before OA-causing injury has taken place^28^. Our previous efforts have demonstrated the improved anti-OA efficacy of liposomal RvD1 compared to free RvD1 in a chow-fed mouse model of PTOA^16^. Hence, we wanted to test our formulations in a more clinically relevant and challenging model of ObOA.

In this study, we show that in a mouse model of ObOA, liposome-based delivery of RvD1 significantly improved joint health than untreated and free RvD1 delivery. Our mechanistic analysis shows that this protective action is mediated via the action of the released RvD1 on cells like macrophages in the synovium.

## Methods

### 1. Materials

The lipids Dipalmitoylphosphatidylcholine (DPPC), 1, 2-Distearoyl-sn-glycero-3-phosphoethanolamine-Poly (ethylene glycol) (DSPE: PEG), Cholesterol, Dioleoyl-3-trimethylammonium propane (DOTAP) were purchased from Avanti polar lipids. Resolvin D1 was purchased from Cayman chemicals. Syringes were purchased from BD Biosciences. Antibodies used: anti-iNOS (NB300-605) and anti-CD206 (NBP1-90020) antibodies were purchased from Novus biological (Centennial, Colorado, USA). Anti-ADAMTS5 (ab41037) and anti-MMP13 (ab39012) antibodies were purchased from Abcam (Cambridge, UK). An anti-β-catenin antibody was purchased from ThermoFisher Scientific (71-2700). Solvents: acetonitrile, methanol, and HPLC grade water were purchased from Fisher Chemicals. All purchased compounds were used without further purification.

### 2. Synthesis and characterization of liposomes

Liposomes were synthesized by the thin-film lipid hydration method. Briefly, the lipids DPPC, DSPE-PEG, and cholesterol were dissolved in chloroform and mixed in their respective molar ratios in a round bottom flask. The chloroform was evaporated using a rotatory evaporator (DLAB RE100 Pro), thus forming thin lipids films. The films generated were hydrated using respective solutions (AF750 in phosphate buffered saline [PBS] for in vivo retention experiments, calcium acetate for all RvD1 loading experiments) at 45°C. The vesicles were then collected and passed through 1 μm and 400 nm membranes to generate liposomes of a defined size. The sizes of liposomes were measured using Malvern Zetasizer μV.

### 3. Cryogenic transmission electron microscopy of liposomes

Liposomes were imaged using a cryogenic transmission electron microscope (Cryo-TEM). Briefly, Holey Carbon Flat R2/2 grids were blotted with a liposomal solution (2 mg/ml) and plunged into liquid ethane using FEI vitrobot to generate vitrified samples. These samples were stored in liquid nitrogen till further use. Furthermore, images of these samples were captured using Thermo Scientific Arctica equipped with a Gatan K2 direct electron detector camera by Latitude S software with a spot size of 7. The total exposure was 40 e^−^/Å2, and the pixel size was 1.2 Å.

### 4. Loading of RvD1 into liposomes

RvD1 was loaded into liposomal using a remote-loading strategy as mentioned earlier^16^. Briefly, thin films of lipids were hydrated with 120 mM Calcium acetate (pH=6) to generate multilamellar vesicles. These vesicles were extruded through filters of different pore sizes (1 μm and 400 nm) to generate liposomes of desired sizes. Finally, the liposomes were pelleted and resuspended in RvD1-containing sodium sulfate solution (pH=4) and loaded at 50 °C for 1.5 h. The loading levels were adjusted to 50 ng RvD1/ mg of lipids. After loading, the formulation was washed twice in PBS and used immediately.

### 5. Release profiles

For characterizing the retention of RvD1, liposomes were incubated in 50% synovial fluid obtained from joints of OA patients (IHEC approval number MSRMC/EC/AP-06/01-2021 at Ramaiah Medical College, Bangalore and 02/31.03.2020 at Indian Institute of Science, Bangalore) for up to 11 days at 37 °C. At each time point, the liposomes were washed and lyophilized. Lyophilized powder was dissolved in 50% isopropanol and the samples were loaded and quantified using HPLC as described elsewhere^16^.

### 6. Mice for *in vivo* studies

The *in vivo* studies were approved by the Institutional Animal Ethics Committee (CAF/Ethics/808/2020). Male C57BL/6 mice (aged 6-8 weeks; weight 20-22g) were used for this study, maintained in individually ventilated cages at the central animal facility (CAF), Indian Institute of Science, Bangalore. Depending upon the group, the mice had *ad libitum* access to either a specialized diet enriched in fat (to model obesity by feeding diet prepared by the National Institute of Nutrition (India) according to Research Diets formulation 12492) or chow (control diet). A cocktail of Ketamine (60 mg/kg) and Xylazine (9 mg/kg) was used to anesthetize animals during procedures.

### 7. Mice model of ObOA

The surgery for modeling ObOA, destabilization of the medial meniscus (DMM) ^28–30^, was performed in mice after 8 weeks of high-fat feeding, as mentioned earlier. The experiments utilized male mice because surgery-induced-OA show more dramatic effects on the cartilage of male mice than that of female mice^31^. One knee joint was subjected to this surgery in each mouse. On the day of the surgery (after completion of 8 weeks of high-fat feeding), the mice were anesthetized using a cocktail of Ketamine (60 mg/kg) and Xylazine (9 mg/kg). After confirming the loss of pedal reflexes, a parapatellar skin incision was made to access the synovium. The synovium was then dissected to expose the underlying joint. The medial meniscotibial ligament (MMTL) was located and surgically transected. After confirming the successful transection of the MMTL by another observer, the synovium and skin were sutured in layers, and a metronidazole wet pack was applied over the sutured area. The mice were then allowed to recover from the anesthesia on a lukewarm surface. Postoperative analgesic care included four subcutaneous doses of Buprenorphine (0.1 mg/kg), once every 12 h. Later, the mice were administered their respective intraarticular treatments over the next three months, as indicated in the result section. The mice were allowed to move freely for the entire duration of the study. During the entire postoperative period, the mice were allowed access to high-fat feed. At the end of the study, the mice were euthanized, and their knee joints were harvested. The joints were fixed in 4% formaldehyde for 6 h and decalcified in 5% formic acid for 5 days. Following decalcification, the joints were dehydrated in multiple gradients of ethanol and xylene and finally embedded in paraffin wax.

### 8. Quantification of serum parameters

Blood was collected from mice using retroorbital venipuncture. Blood was then allowed to clot at room temperature for 2 h, following which the collection tubes were centrifuged (8000g, 10 mins, 4°C). In this process, the clotted blood forms a pellet at the bottom of the tube, while the liquid fraction (serum) of the blood is at the top. From these tubes, the supernatants were collected and used for further analysis. Analysis kits for total cholesterol, LDL-cholesterol, and total triglycerides were quantified using colorimetric kits from Delta Lab (Maharashtra, India) as per the manufacturer’s instructions.

### 9. Histology and staining

The tissues embedded in paraffin blocks were sectioned into 5 μm thick sections using Leica HistoCore MULTICUT and collected on poly-L-lysine-coated glass slides. The sections were hydrated using a series of ethanol gradients and stained using the Safranin-O^32^. The severity of the disease was quantified on a scale of 0-24 using a scoring protocol prescribed by OARSI ^33^. This protocol considers the joint’s damage, including loss of proteoglycan, chondrocyte apoptosis, and the presence of osteophytes^33^. In this study, the medial surfaces were used for analysis because lateral surfaces did not show significant damage (**Figure S1**). Scoring was done by trained veterinarians blinded to the study.

### 10. Measurement of synovial thickness

To quantify fibrosis, we measured the thickness of the synovial membrane. Briefly, joint sections were stained with the safranin-O staining technique (as mentioned earlier) and images of the respective synovial membranes were captured. The thicknesses of the synovial membranes covering the medial regions of the joints were then measured at 3 different locations along their lengths using ImageJ.

### 11. Quantification of cells in the synovial membrane

We quantified the severity of synovitis by measuring cellular density in the synovial membrane. Briefly, joint sections were stained with the safranin-O staining technique (as mentioned earlier) and images of the respective synovial membranes (~3 images from each synovial membrane) were captured. The number of cells was counted using specialized macros implemented in ImageJ. This number was normalized to the area of the tissue used to quantify the number of cells.

### 12. Immunohistochemistry

IHC was performed to quantify iNOS^+^ cells, CD206^+^ cells, MMP13^+^, and ADAMTS5^+^ regions in the synovium and cartilage. This method of IHC was derived from our previous studies where several other markers like iNOS, CD206, and LC3B were used for cellular analysis^16, 34^. Briefly, heat-induced epitope retrieval was performed using Tris-EDTA overnight treatment at 65°C, followed by retrieval with 1N HCl and Trypsin-CaCl2 (0.05% Trypsin in 1xPBS and 1% CaCl2). The resulting sections were then incubated with primary antibody (concentrations of antibodies used: anti-iNOS-1 μg/mL; anti-CD206-0.66 μg/mL; anti-MMP13-1 μg/mL; anti-ADAMTS5-0.33 μg/mL) for 16 h. These sections were then washed to remove excess unbound antibody and incubated with horseradish peroxidase (HRP)-conjugated secondary antibody (1.6 μg/mL)for 2 h. After washing the unbound secondary antibody, sections were incubated with 3,3’Diaminobenzidine (DAB) substrate for 1 h. Excess unreacted DAB was washed, and images were captured using an Olympus BX53F brightfield microscope. The images were thresholded against the background and counted using the ‘analyze particles’ module in ImageJ.

### 13. Testing for mechanical allodynia

OA-associated allodynia was tested using von Frey filaments (Aesthesio tactile sensory filament, Ugo Basile). Briefly, the animals were introduced in a customized cage with perforated bottom and were allowed to be acclimated to it for a few minutes. A von Frey filament representing the smallest force was pushed against the plantar regions of the paw of respective animals through the perforations in the cage. This exercise was performed ten times per filament. If the animal withdrew its paw three out of ten times, the force represented by that filament was noted as the paw withdrawal threshold. If no such activity was seen, the exercise was continued with the next filament. Only animals with no visible injury were used for data collection.

### 14. Statistics

Data presented in this manuscript were represented as mean ± standard deviation with at least 3 replicates in each group unless stated otherwise. Data were analyzed using one-way ANOVA for normal distributions and using other non-parametric tests (e.g., Dunn’s or Kruskal-Wallis tests) for ordinal datasets. Outliers were analyzed using the Grubbs test (α=0.05). If an outlier was detected in any assay, then the data from that animal was completely excluded from all other assays. The 95% confidence interval was considered significant. Power analysis (power=0.8, α=0.05) yielded the desired sample size of 8-11 mice per group.

## Results

### 1. Lipo-RvD1 creates a depot of RvD1 in human synovial fluid

Liposomes were synthesized using the thin film hydration followed by extrusion through 400 nm pore size membranes as described earlier^16^. The size and zeta potential was found to be 312±10.1 nm and −1.92±0.05 mV respectively (**figure 1a, 1b**). Synthesis was followed by loading of RvD1 into liposomes using an active encapsulation strategy (E.E.=55.6±24.5%). To evaluate the sustained release of RvD1 from the formulation in the synovium-like conditions, lipo-RvD1 was incubated in synovial fluid (obtained from patients undergoing joint replacement surgery) at 37°C for various time intervals. At each of these intervals, liposomes were collected and the drug retained was quantified using HPLC. HPLC quantification showed that RvD1 molecules were retained intraliposomally for ~11 days (**figure S2**).

**Figure 1.**
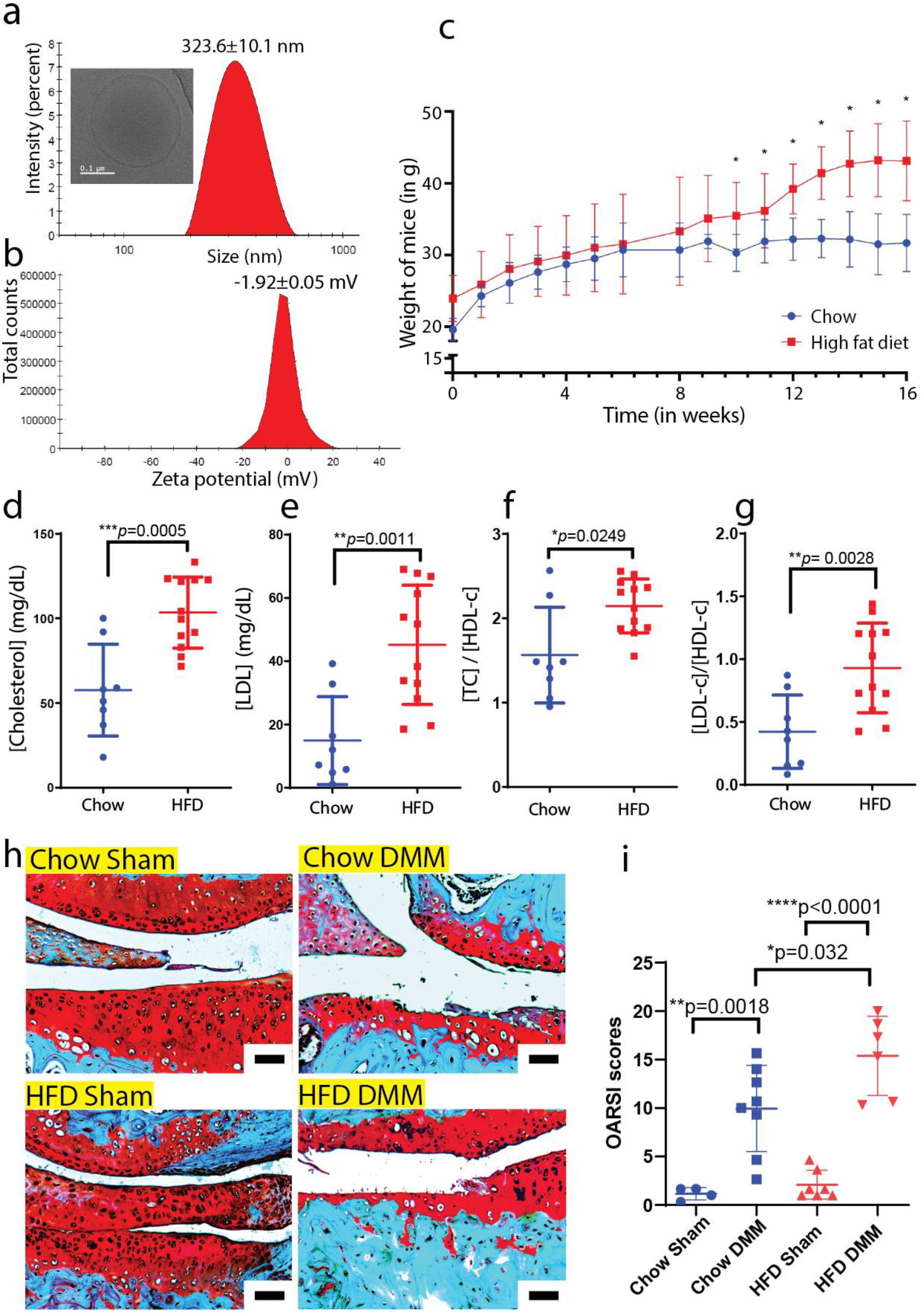
High-fat fed mice become obese and show severe cartilage damage after DMM surgery. **(a)** Size estimation of liposomes using DLS and cryoTEM (inset). **(b)** Surface charge of liposomes as indicated using zeta potential. **(c)** Body weight profiles of mice fed high-fat diet (HFD) and chow. Serum levels of **(d)** cholesterol, **(e)** LDL, and the serum ratios **(f)** TC/HDL-c and **(g)** LDL-c/HDL-c; n=8 lean animals and n=12 obese animals. **(h)** Characteristic safranin-O stained sections of cartilage (scale bar 50 μm). **(i)** OARSI scores indicating the severity of the joint damage; n=4-8 animals per group. For **d**, **e**, **f**, **g**, and **i**, \**p*<0.05, \*\**p*<0.01, \*\*\**p*<0.001, and \*\*\*\**p*<0.0001 between the respective groups indicated in the figures using ANOVA followed by Tukey’s posthoc test. For **c,** \*\**p*<0.01 between the respective groups indicated in the figures using multiple t-tests. Values are expressed as mean ± SD. Scale bar 50 μm.

### 2. Obesity can be induced in mice by high-fat feeding

Obesity is characterized by an increase in body weight due to the deposition of excessive fat in adipose tissue^35^. This disorder can be modeled in mice by providing *ad libitum* access to specialized diets which are enriched in fat^36, 37^. In our experiments, we used a specialized feed containing 60% fat by calorie content to generate overweight mice. We observed a statistically significant difference between the weights of the chow-fed and HFD-fed mice from the 8^th^ week after the commencement of feeding (**figure 1c**). Obese patients often show systemic dyslipidemia and upregulated low-density lipoprotein fraction of cholesterol (LDL-c), triglycerides, and cholesterol ^38, 39^. Accordingly, we found that the total cholesterol (TC) was higher in the serum of overweight mice than in their leaner counterparts (**figure 1d**). LDL-c also increased with dietary fat in mice (**figure 1d**) and the ratio of both TC (**figure 1f**) and LDL-c (**figure g**) to the high-density lipoprotein fraction of cholesterol (HDL-c) was higher in overweight mice than in their leaner counterparts. Since these parameters were in alignment with those reported in the literature^40^, high fat-fed mice were successfully classified as obese.

The destabilization of the medial meniscus (DMM) is a widely used surgical model for obesity-related post-traumatic OA (PTOA)^28, 30^. The medial meniscus, located between articulating surfaces, provides stability to the joint by absorbing mechanical shock. Surgically cutting and dislocating this tissue mimics damage resulting from traumatic events like sports injuries and accidents and results in contact between the two articulating surfaces. This surgery generates OA-like changes over 1-3 months. To mimic ObOA, we performed DMM in obese mice and compared it with DMM in mice fed with standard diet (11% kCal by fat). We found more severe damage to the cartilage in obese mice than in their leaner counterparts after DMM surgery (*p*=0.032) (**figure 1h, 1i**). Furthermore, we observed that obesity alone was not sufficient to increase the severity of OA, as seen from the comparison between the pathologies of lean and obese sham joints (**figure 1h, 1i**). We also showed that the articulating surfaces in both the chow-fed DMM and high-fat diet DMM were severely denuded, with more damage expressed in the latter (damage in obese mice is ~1.5x more severe compared to their leaner counterparts). This finding is in agreement with earlier studies which showed that obesity is the major contributor to the pathology^7^.

### 3. Lipo-RvD1 is a good prophylactic candidate for ObOA treatment

We performed DMM surgery in obese mice and used a prophylactic dosing regimen by injecting freshly synthesized lipo-RvD1 intraarticularly (IA) at weeks 1 and 4 after surgery (**figure 2a**). Lipo-RvD1 arrested the progressing cartilage damage and maintained the overall joint integrity. In the case of lipo-RvD1-treated mice, the OARSI scores were similar to sham mice. There was a 6-8-fold reduction in OARSI scores of lipo-RvD1 treated mice compared to DMM joints (*p*<0.0001), which had complete loss of articulating cartilage at certain sites of damage (**figure 2b, 2c**). IA injection of blank liposomes failed to show any efficacy (**figure S3**). **Figures 2b**, and **2c** also depict that more chondrocytes were observed to be healthy in the lipo-RvD1 treated joint than in free RvD1 and DMM joints. Administration of free RvD1 also showed some protective effect on the joint but the OARSI scores were still significantly higher than sham group (*p*<0.0001). Several inflammatory diseases have an imbalance between the M1 and M2 macrophages^30, 41, 42^. The ratio of M1/M2 cells is skewed in OA as well and proinflammatory cytokines from M1 macrophages drive cartilage damage^43^. Previously it was shown that prophylactically administered RvD1 improves the ratio of M1/M2 macrophages in the synovial membrane^16, 28^. As seen from our results, DMM mice had higher levels of iNOS^+^ M1 cells than sham mice, which indicates the presence of a proinflammatory environment in the synovium (*p*=0.0072) (**figure 2d, 2e**). Furthermore, lipo-RvD1 treatment promoted polarization towards M2 cells as compared to all other groups, evident by an increased number of CD206^+^ cells (**figure 2f, 2g**). Overall, we observed that lipo-RvD1 formulation was increasing M2 macrophages in the joint which reduced the net inflammatory activity within the synovium and promoted clearance of debris and other inflammatory factors. This inference, however, could not be extrapolated to other regions of the joint(**figure S4**). In OA, chondrocytes release catabolic enzymes like ADAMTS5 and MMP13, which also are recognized as markers of chondrocyte hypertrophy^44^. Administration of lipo-RvD1 suppressed the expression of the catabolic mediators MMP13 (but not ADAMTS5), thus preventing the formation of hypertrophic chondrocytes in mice (**figure S5**).

**Figure 2.**
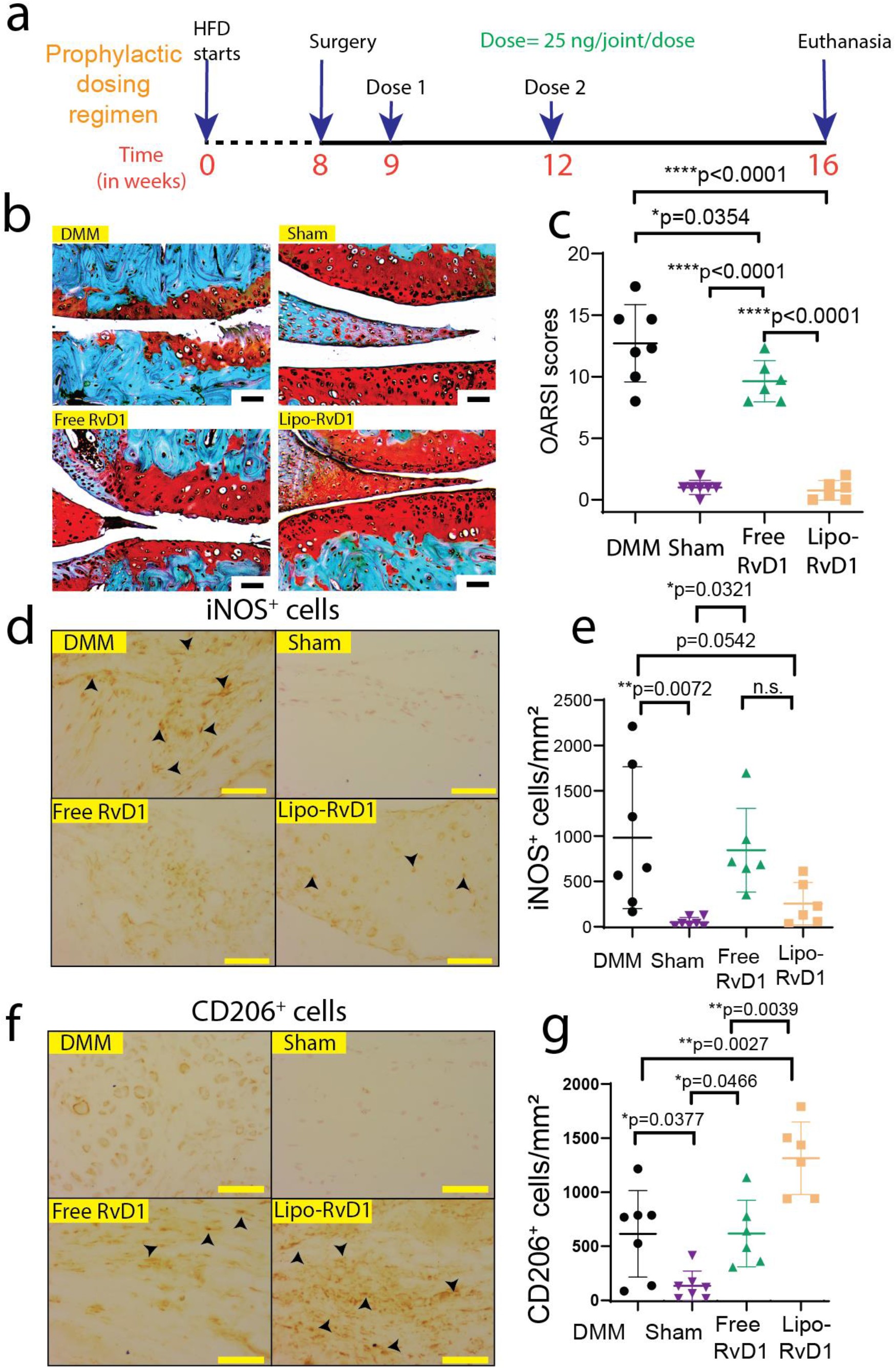
Prophylactic administration of lipo-RvD1 maintains cartilage integrity. (**a**) Timeline of the experiment. (**b**) Characteristic Safranin-O-stained histology sections of different groups of mice (scale bar 200 μm). (**c**) OARSI scores of Safranin O-stained sections of mice knee joints administered with respective treatment; n=7-8 animals per group. (**d**) IHC images depicting levels of iNOS^+^ M1 macrophages synovial membrane (scale bar 50 μm). (**e**) Quantification of iNOS^+^ M1 macrophages in the synovial membrane; n=7-8 animals per group. (**f**) IHC images depicting levels of CD206^+^ M2 macrophages in synovial membrane (scale bar 50 μm). (**g**) Quantification of CD206^+^ M2 macrophages in the synovial membrane; n=7-8 animals per group. For **c**, **e**, and **g** two points in the free RvD1 and two in lipo-RvD1 group were removed after outlier analysis (Grubbs test). For **c**, **e,** and **g** \**p*<0.05, \*\**p*<0.01, and \*\*\*\**p*<0.0001 between the respective groups indicated in the figures using ANOVA followed by Tukey’s posthoc test. Values are expressed as mean ± SD. Scale bar 50 μm.

### 4. Lipo-RvD1 is effective as therapeutic treatment in ObOA

In clinics, the diagnosis of OA relies on radiographic evidence of joint damage, which is visible only when the damage has progressed significantly^45^. Accordingly, therapeutic formulations are critical for the successful treatment of OA. Hence, we tested the therapeutic efficacy of lipo-RvD1 by injecting it at 3 and 6 weeks after DMM surgery. This timeline was chosen as a suitable therapeutic regimen because cartilage damage is known to start within two weeks after DMM^46^ (**figure 3a**). In this challenging regimen, IA-administered lipo-RvD1 significantly reduced OARSI scores compared to free RvD1-administered joints (*p*=0.0006) and DMM-only joints (*p*=0.0001) (**figure 3b, 3c**). M1 macrophages were present in large numbers in the synovial membrane of the DMM-treated joints compared to the lipo-RvD1-treated joints, but the difference was not statistically significant (**figure 3d, 3e**). However, the therapeutic regimen of administration of lipo-RvD1 showed increased levels of pro-resolution M2 macrophages compared to DMM (*p*=0.0002) or free RvD1 (*p*=0.0006) (**figure 3f, 3g**). We further observed that the lipo-RvD1 treatment reduced the expression of damaging enzymes like MMP13 and ADAMTS5 compared to free-RvD1 treatment and protected cartilage from degradation (**figure S6**). To evaluate the minimum required doses of lipo-RvD1 to achieve anti-OA efficacy, we further reduced and administered one dose (given at 12 weeks) over the total experiment (**figure S7a**). Our OARSI scores and histology results showed that this regimen was not sufficient to protect the joint from impending OA-related joint damage (**figure S7b-c**). However, it is possible that administration of lipo-RvD1 can improve efficacy if given at an earlier time point which was not explored in this study.

**Figure 3.**
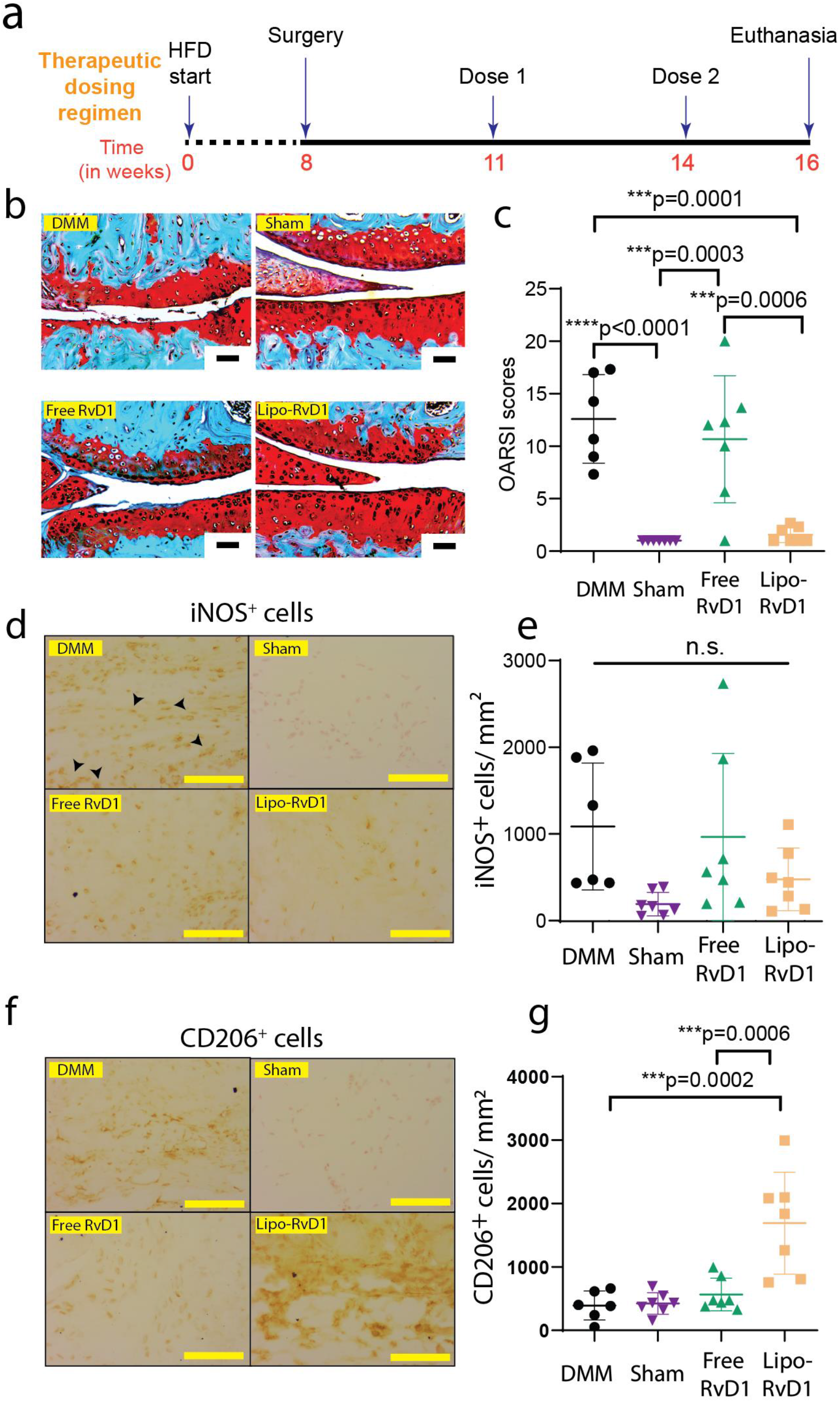
Therapeutic administration of lipo-RvD1 protects cartilage from progressing damage. (**a**) Timeline for the study. (**b**) Safranin-O-stained characteristic histological images of different groups of animals (scale bar 200 μm). (**c**) OARSI scores of Safranin O-stained sections of mice joints administered with respective treatment; n=6-8 animals per group. (**d**) Characteristic IHC images depicting levels of iNOS^+^ M1 macrophages in the synovial membrane of respective mice joints (scale bar 50 μm). (**e**) Quantification of iNOS^+^ M1 macrophages in the synovial membrane of respective mice joints; n=6-8 animals per group. (**f**) IHC images depicting levels of CD206^+^ M2 macrophages in synovial membrane (scale bar 50 μm). (**g**) Quantification of CD206^+^ M2 macrophages in the synovial membrane; n=6-8 animals per group. For **c**, **e**, and **g**, \*\*\**p*<0.001 and \*\*\*\**p*<0.0001 between the respective groups indicated in the figures using ANOVA followed by Tukey’s posthoc test. n.s. = non-significant. Values are expressed as mean ± SD. Scale bar 50 μm.

Wnt signaling plays a major role in OA pathology^47^ and inhibitors of this pathway (e.g. lorecivivint-phase 3 clinical trials; NCT04931667) are currently in clinical trials as an anti-OA medication^48^. β-catenin is a mediator of Wnt signaling and is upregulated in OA^47^. In our prophylactic study, we observed that both free RvD1 and lipo-RvD1 administration to OA joints suppressed the overexpression of β-catenin in chondrocytes (**figure S8a**), thus indicating the ability of RvD1 to regulate Wnt signaling. Furthermore, in our therapeutic study, we observed that administration of free RvD1 was not sufficient to prevent overexpression of β-catenin, and this regulation of Wnt signaling was achieved only if RvD1 was administered in a sustained manner through liposome-encapsulated RvD1 (**figure S8b**). It is not clear how RvD1 administration is modulating Wnt signaling and more experiments are required to understand this phenomenon.

One of the hallmarks of advanced OA is the development of chondrophytes, which develop into osteophytes. These characteristics are also observed in the DMM model for OA^16^. In our studies, although some of the joints demonstrated these features, a majority of them were devoid of chondrophytes (**figure S9**).

### 5. Lipo-RvD1 reduces synovitis in mice

Synovitis is a hallmark of OA^49^ and is characterized by an increased pool of leukocytes in the synovial membrane. Synovitis increases the production of inflammatory mediators which leads to chondrocyte hypertrophy and blocks anabolism^50^. Our data showed that treating DMM mice with lipo-RvD1 reduced the synovial membrane (SM) thickness (**figure 4a, b, c, d**) and cellularity (**figure 4e, f, g, h**) as compared to DMM-only mice, in both, therapeutic and prophylactic regimens of administration. SM from lipo-RvD1-treated joints was similar in thickness to SM in sham joints in both the therapeutic and prophylactic regimens of administration, thus emphasizing the ability of lipo-RvD1 to suppress excessive fibrosis of SM (**figure 4a, b, c, d**). We observed that in the therapeutic regimen, lipo-RvD1 treated membranes were thinner than free RvD1 treated membranes (*p*=0.0118). This result also held true for the total cellularity of the SM (**figure 4e, f, g, h**). Lipo-RvD1 treated joints showed a better ability to prevent cells from infiltrating the inflammatory milieu compared to free RvD1 when administered therapeutically (*p*=0.0002) (**figure 4g, h**).

**Figure 4:**
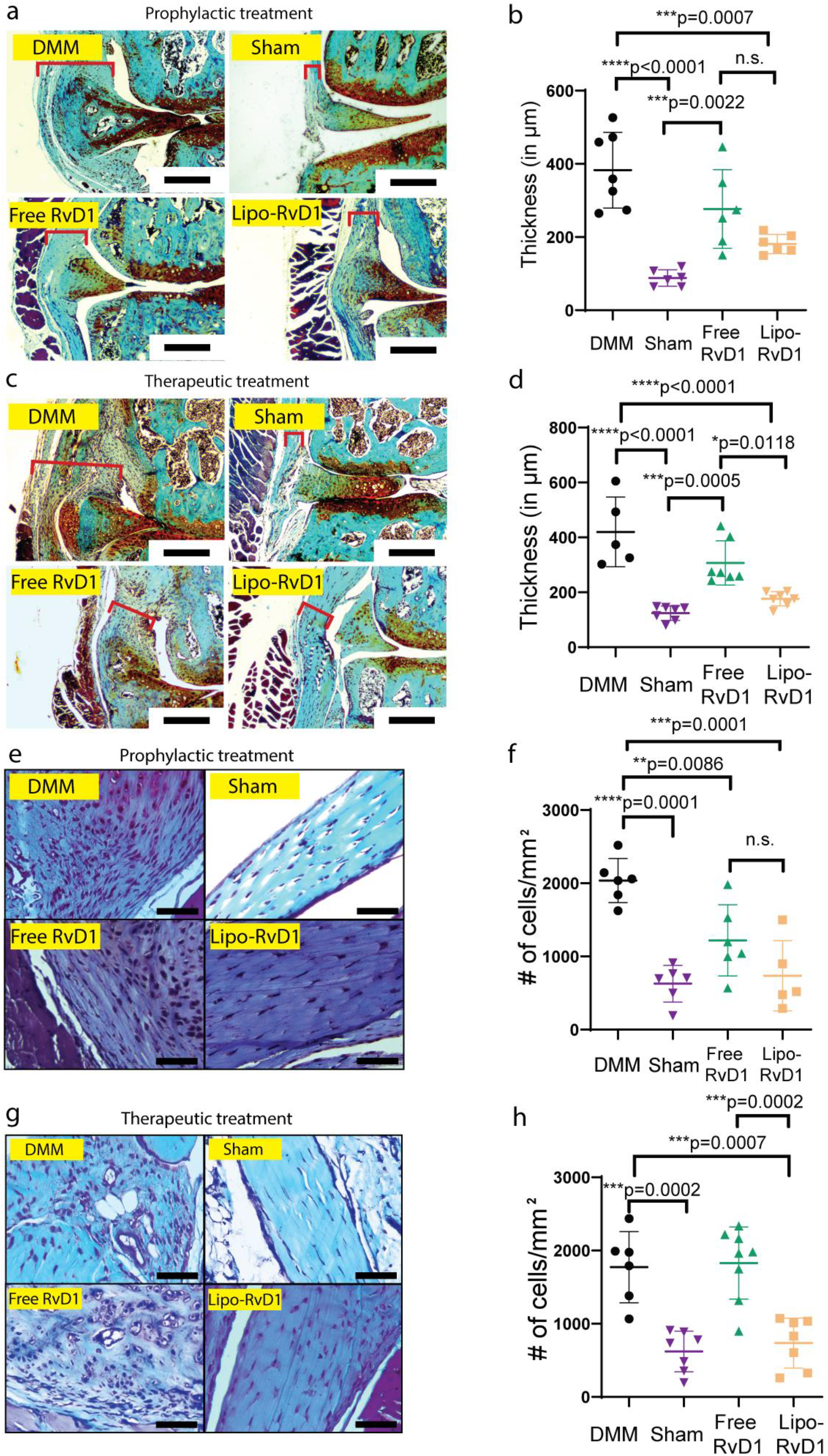
Lipo-RvD1 treated mice show reduced synovitis. (**a**) Images of stained sections of synovial membranes of joints treated with lipo-RvD1 prophylactically. (**b**) Thickness of the synovial membrane of joints treated with lipo-RvD1 prophylactically; n=6-8 animals per group. (**c**) Images of stained sections of synovial membranes of joints treated with lipo-RvD1 therapeutically. (**d**) Thickness of the synovial membrane of joints treated with lipo-RvD1 therapeutically; n=5-8 animals per group. (**e**) Magnified images of stained sections of synovial membranes of joints treated with lipo-RvD1 prophylactically. (**f**) Quantification of cells in the synovial membrane of joints treated with lipo-RvD1 prophylactically; n=6-8 animals per group. (**g**) Magnified images of stained sections of synovial membranes of joints treated with lipo-RvD1 therapeutically. (**h**) Quantification of cells in the synovial membrane of joints treated with lipo-RvD1 therapeutically; n=6-8 animals per group. For **b**, **d**, **f**, **h,** \**p*<0.05, \*\**p*<0.01, \*\*\**p*<0.0001, and \*\*\*\**p*<0.0001 between the respective groups indicated in the figures using ANOVA followed by Tukey’s posthoc test. n.s. = non-significant. Values are expressed as mean ± SD. Scale bar 50 μm.

### 6. Lipo-RvD1 reduces the incidence of OA-associated allodynia

OA patients suffer from pathological pain (allodynia)^51^. Pain threshold was tested using Von Frey filaments 8 weeks after surgery (immediately before euthanasia). We observed that administration of lipo-RvD1 was more effective in alleviating the allodynia than free RvD1 injected mice (*p=0.0009*) in the prophylactic regimen of administration (**figure 5a**) but not in the therapeutic regimen (**figure 5b**). IA administration of free RvD1 did not generate adequate analgesia in the therapeutic regimen, and the paw-withdrawal threshold of these mice was not statistically different than that of DMM-operated mice (**figure 5b**). Our results from the therapeutic regimen showed that lipo-RvD1 improves the pain threshold of the limb, but the difference was not statistically significant compared to free RvD1(**figure 5b**).

**Figure 5:**
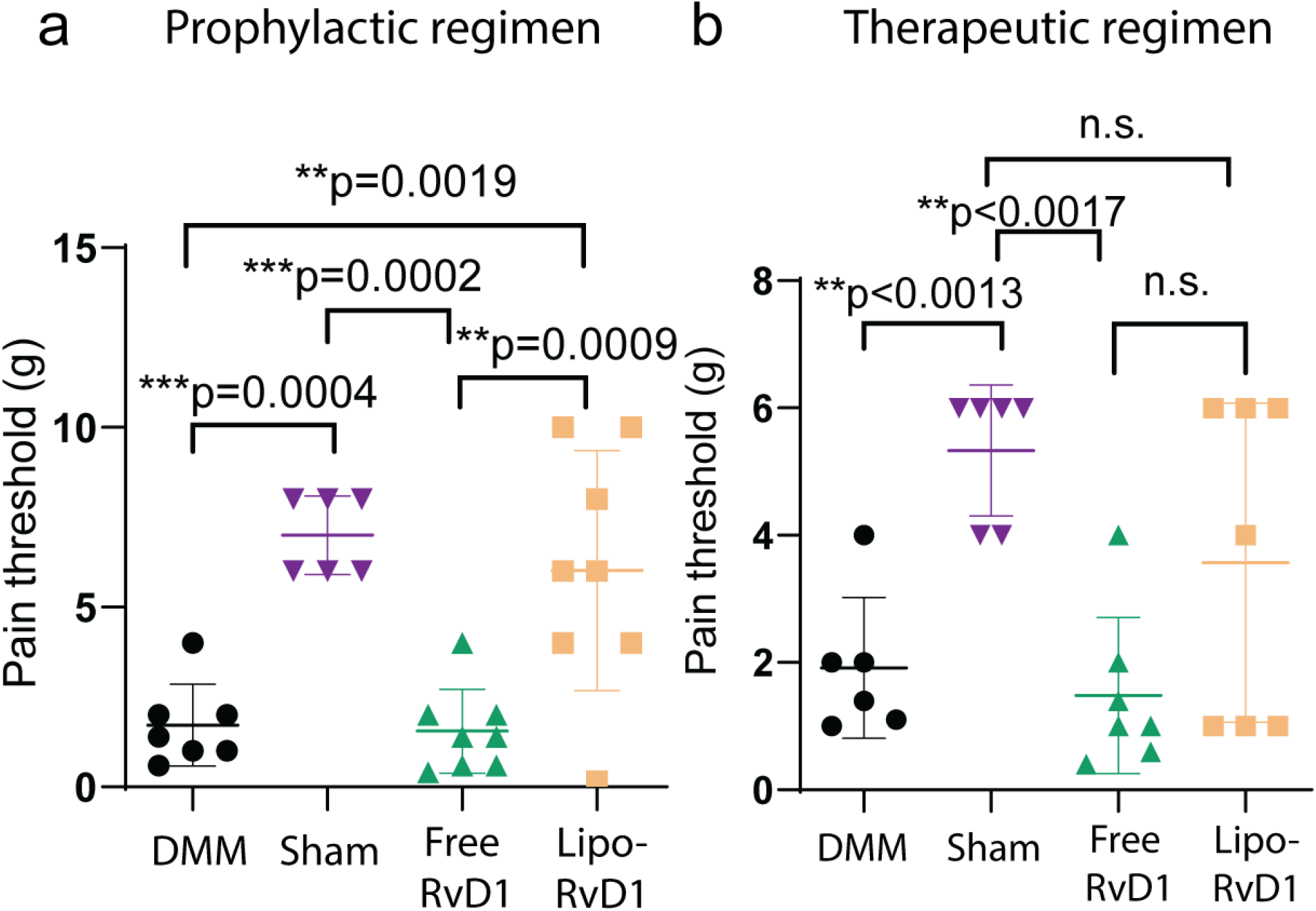
Lipo-RvD1 shows analgesia in prophylactic but not in the therapeutic regimen. Paw withdrawal thresholds were obtained using von Frey filaments in mice administered lipo-RvD1 (**a**) prophylactically and (**b**) therapeutically; n=6-8 mice per group in both studies. For **a** and **b**, \*\**p*<0.01, \*\*\**p*<0.001, and \*\*\*\**p*<0.0001 between the respective groups indicated in the figures using ANOVA followed by Tukey’s posthoc test. n.s. = non-significant. Values are expressed as mean ± SD. Scale bar 50 μm.

## Discussion

Obesity is one of the most common sources of metabolic dysfunction in humans and contributes to the pathology of several diseases by mediating metabolic dysregulation, including OA^3, 52^. In obesity, the adipose tissue is affected, which acts as an endocrine organ to release several proinflammatory adipokines and cytokines like leptin^6^ and IL-6^8^. The ratio of ω-3 to ω-6 fatty acids is also skewed in the adipose tissue^53^. Studies have shown that RvD1 is downregulated in obese tissue^54^. Improving this ratio by increasing the dietary ω-3 fatty acids is known to improve the inflammatory tone of the tissue^55,56^ (this phenomenon is also observed in OA^57, 58^). Reports also suggest that RvD1 (an ω-3 fatty acid) reduces the levels of leptin in adipocytes *in vitro^59^*, thus promoting healthy adipose tissue. We had previously shown that administration of lipo-RvD1 was efficacious in chow-fed mice^16^. All these studies motivated us to explore the effect of RvD1 in the treatment of OA in obese mice.

Macrophages are immune cells with significant plasticity which allows them to perform diverse roles like wound healing, tissue homeostasis, and protection against inflammation^60^. Traditionally, the entire spectrum of macrophages could be divided into two broad phenotypes-M1 phenotype, which is responsible majorly for a proinflammatory response, and M2 macrophages, which are dominant in a proresolution medium^60^. These phenotypes are dynamic and can change depending on the factors present in the surrounding. In the case of several inflammatory diseases, chronically activated proinflammatory M1 macrophages are the major aggressors of tissue damage, and increasing the fraction of M2 macrophages is a viable strategy for the treatment of such diseases^28, 60^. We observed that administration of lipo-RvD1 improves the levels of M2 macrophages while simultaneously suppressing the M1 phenotype compared to DMM joints. Furthermore, for the dose of RvD1 that we administered (25 ng/ joint), this phenomenon was observed only when the molecule was administered in a liposome-encapsulated, controlled-release format. We did not observe a significant reduction of M1 macrophages in the synovial membrane of lipo-RvD1 treated joints compared to free-RvD1. In this study, we used iNOS and CD206 as markers for M1 and M2 macrophages respectively. While these markers are well-accepted^28, 61^, endothelial cells and other immune cells also express CD206 and iNOS respective. Other markers of macrophages and its subtypes can further increase confidence in data.

Liposomes have several advantages like biocompatibility, flexible synthesis, and stability for a long duration. For these reasons, liposomes are the most commonly approved sustained-release formulations for human use. In our previous study, we also reported that liposome-mediated delivery of drugs improved their IA retention compared to free molecules^16^.

Pathological neuropathic pain is a major consequence of OA, and has two common symptoms: allodynia(pain due to stimuli that is not normally painful) and hyperalgesia (increased sensitivity to painful stimuli)^62^. The nature of pain depends on the nature of the stimuli generating the pain^62^. This study evaluates allodynia (pain generated from von Frey filaments that are normally considered painless), but does not investigate the hyperalgesic behavior of mice in the respective group. This limitation prevents us from gaining a broad perspective on the analgesic properties of Lipo-RvD1 and will be addressed in future experiments.

The Transient Receptor Potential (TRP) family of mediators is critical in the response to mechanical stimuli, including those inducing pain^63^. In OA, members of this family of receptors, especially TRPV1 and TRPV4, are associated with the severity of pain^64–66^. RvD1 has been shown to generate analgesia in different models by targeting a few members of this family, like TRPV3, TRPV4, and TRPA1^67, 68^. RvD1 receptors are also known to play a role in analgesia^69^. Our results show that the sustained presence of RvD1 in the affected knee joint helps alleviate OA-associated pain, especially under a prophylactic regimen. The ability of lipo-RvD1 to provide analgesia can ensure higher patient compliance and also improve the translational and clinical relevance of the formulation. Several other mechanisms of pain have been discovered, including the role of mechanotransducers like PIEZO^70^. However, no direct interaction between RvD1 and these receptors have been discovered, and hence their role in the proresolution activity of RvD1 remains elusive.

One of our interesting findings includes the ability of lipo-RvD1 to provide analgesia in prophylaxis, but not when administered as a therapeutic agent. In OA (both, human OA^71^ and collagen-induced mouse models of OA^72^), cartilage lesions and damaged nociceptors lead to neuropathic pain. In this study, we hypothesize that prophylactic administration of RvD1 prevents the initial surge in inflammatory factors, thus effectively inhibiting tissue damage. On the contrary, as the therapeutic administration is done later, it is possible that excessive early inflammation causes nerve damage and subsequent neuropathic pain to prevail.

Our administration frequencies were inspired by Sun *et.al*., with certain changes^28^. The first (of the two injections) administration for prophylactic treatment was performed 5-6 days after the surgery as such a prophylactic dose is feasible for human use in case of severe trauma to the joints. Similarly, the first dose for the therapeutic regimen was administered three weeks after surgery because, by this time, the pathology in the joint is reported to progress significantly^30, 46^.

Recent reports show that body composition, not body weight, has a true bearing on joint health^73–75^. Although we show that an increase in body weight by sustained high-fat feeding increases serum lipids (an indicator of high body fat), this remains indirect evidence of a body composition comprised of high fat. In future experiments, direct quantification of body composition can be performed using powerful techniques like DEXA scanning^76^.

A limitation of this study is that the experiments in this study utilized only male mice as surgery-induced-OA shows more dramatic effects on the cartilage of male mice than that of female mice^31^. ObOA can be reproducibly mimicked in HFD-fed obese mice by performing DMM surgery on the respective stifle joints. This model also shows certain similarities with the etiology of disease in humans^77^. However, it is not possible to mimic the load-bearing patterns in the articulating joints of large bipeds (humans) in small quadrupeds like mice^78, 79^. Similarly, the vast difference in the sizes of the joints results in unpredictable and unscalable pharmacokinetics of the molecule, which accounts for poor translation to clinics. Hence for future translation, further testing is required in larger animals. Another limitation of this study is the inability to detect osteophytes using the techniques described earlier. This detection can be performed in future studies using more sensitive techniques like microCT.

## Conclusion

In this study, we formulated nanoliposomes that arrested cartilage damage in an ObOA model of mice. The controlled release of RvD1 was achieved for ~11 days. Lipo-RvD1 promoted the M2 polarization of macrophages in the synovium which mediated resolution of inflammation resulting in substantially less cartilage damage. This formulation also reduced disease symptoms like OA-associated pain when used as a prophylactic regimen. Lipo-RvD1 has shown encouraging efficacy as both a prophylactic and therapeutic agent and can thus be a promising strategy to treat OA.

## Ethics statement

All *in vivo* experiments were approved by the Institute Animal Ethics Committee (CAF/Ethics/808/2020). Experiments that utilized synovial fluid were approved by the Institutional Human Ethics committee (IHEC) (02/31.03.2020) and MS Ramaiah Medical College, Ethics Committee (MSRMC/EC/AP-06/01-2021).

## Acknowledgments

We thank Prof. Sathees Raghavan and Prof. Sanhita Sinharay for access to the instruments used in this study. We also thank Central Animal Facility (CAF) for breeding and maintaining mice. Dr. Dhanusha G and Dr. Thirumala M’s help with scoring histological sections is also acknowledged. Finally, we would like to acknowledge Early Career Research Award (Science and Engineering Research Board, Department of Science and Technology, India, **ECR/2017/002178**), Har Gobind Khorana Innovative Young Biotechnologist Award (Department of Biotechnology, **BT/12/IYBA/2019/04**), Indian Institute of Science start-up grant, Private funding from Mr. Lakshmi Narayan, and Private funding from Dr. Vijaya and Rajagopal Rao funding for Biomedical Engineering research at the Centre for BioSystems Science and Engineering and express our gratitude for funding this research. The authors do not declare any conflicting interests.

