## Supplementary Information for "Sustained release Resolvin D1 liposomes are effective in the treatment of osteoarthritis in obese mice"

***Corresponding author**

**Supplementary Information**

`

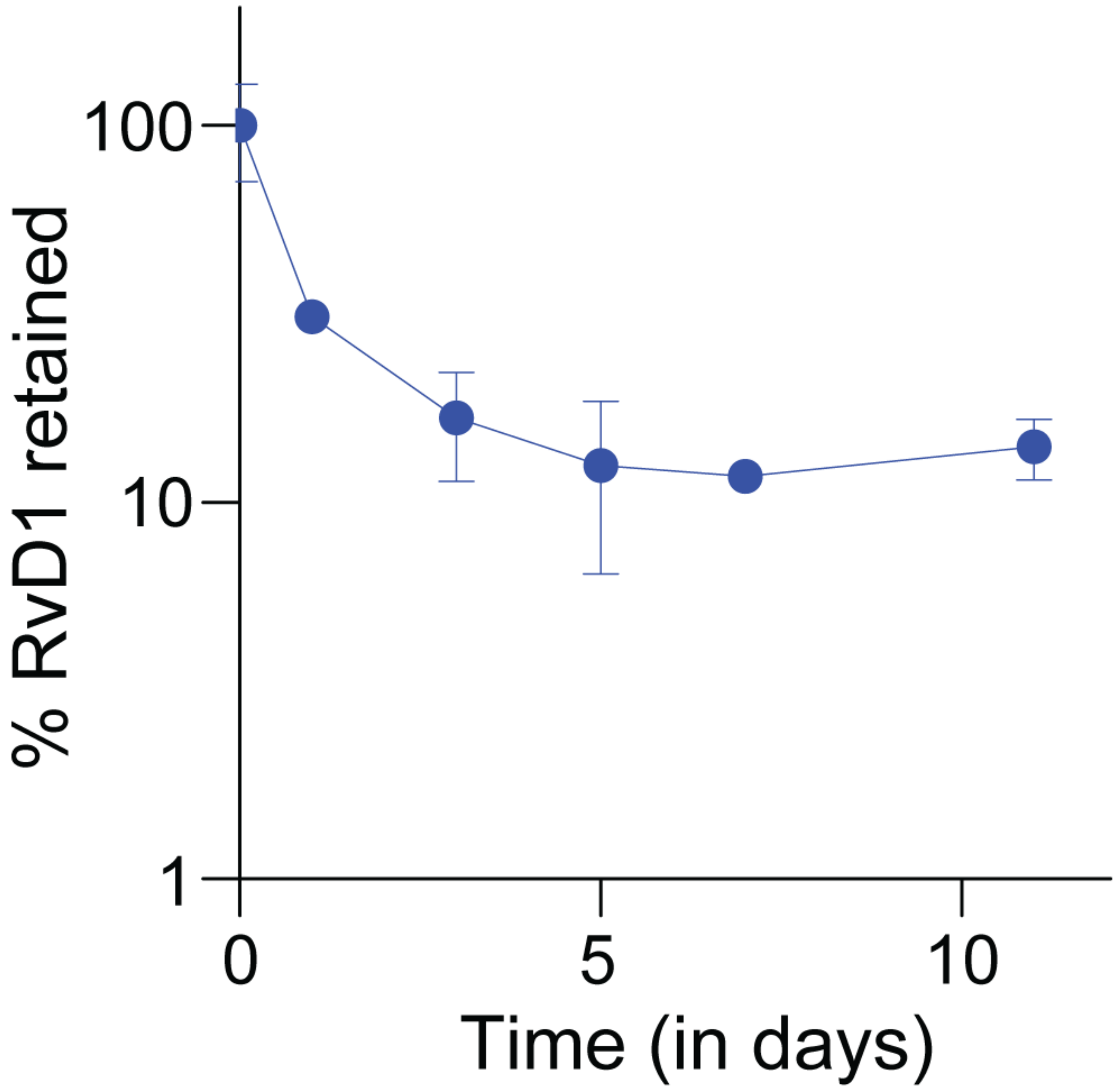


**Figure S1. Lipo-RvD1 shows sustained release of RvD1 in synovial fluid.** Release profile of lipo-RvD1 in synovial fluid obtained from OA patients. n=3-5 replicates per time point.

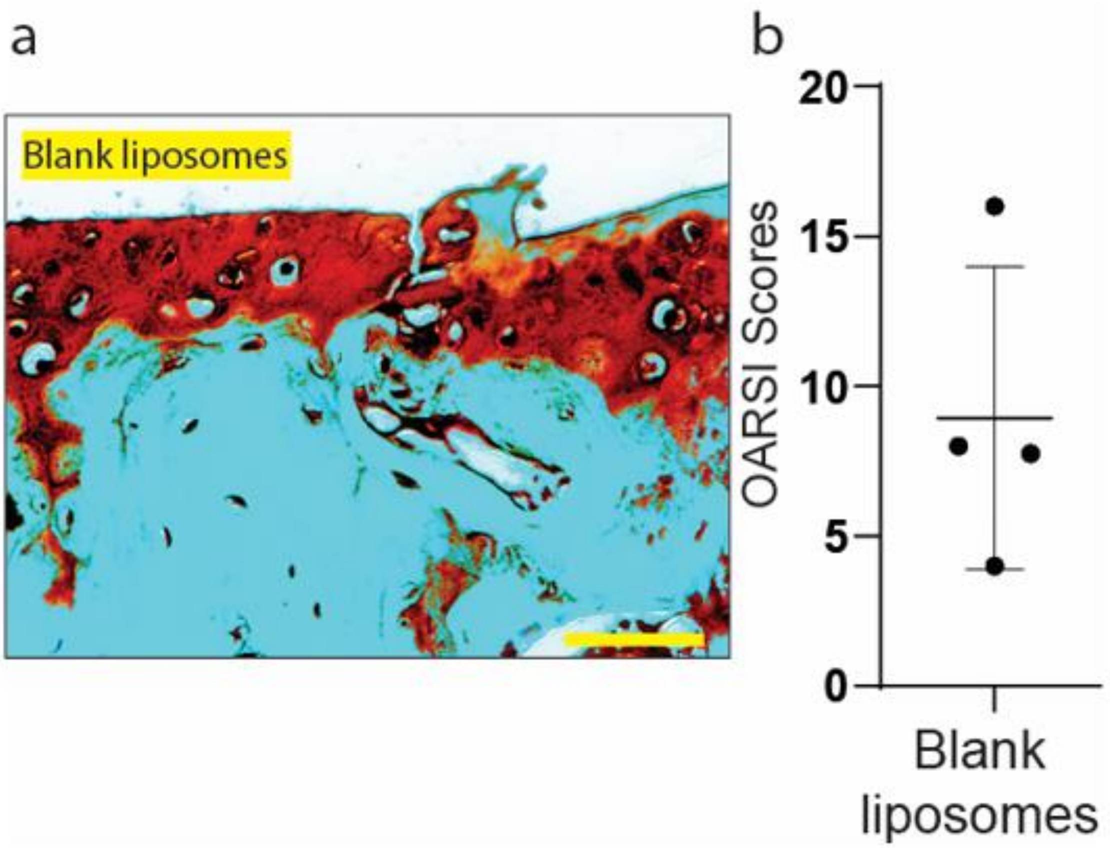


**Figure S2: IA injection of blank liposomes does not reduce the severity of OA.** (**a**) Safranin O-stained sections and (**b**) OARSI scores of joints treated with blank liposomes; n=4 joints. Scale bar 50 µm.

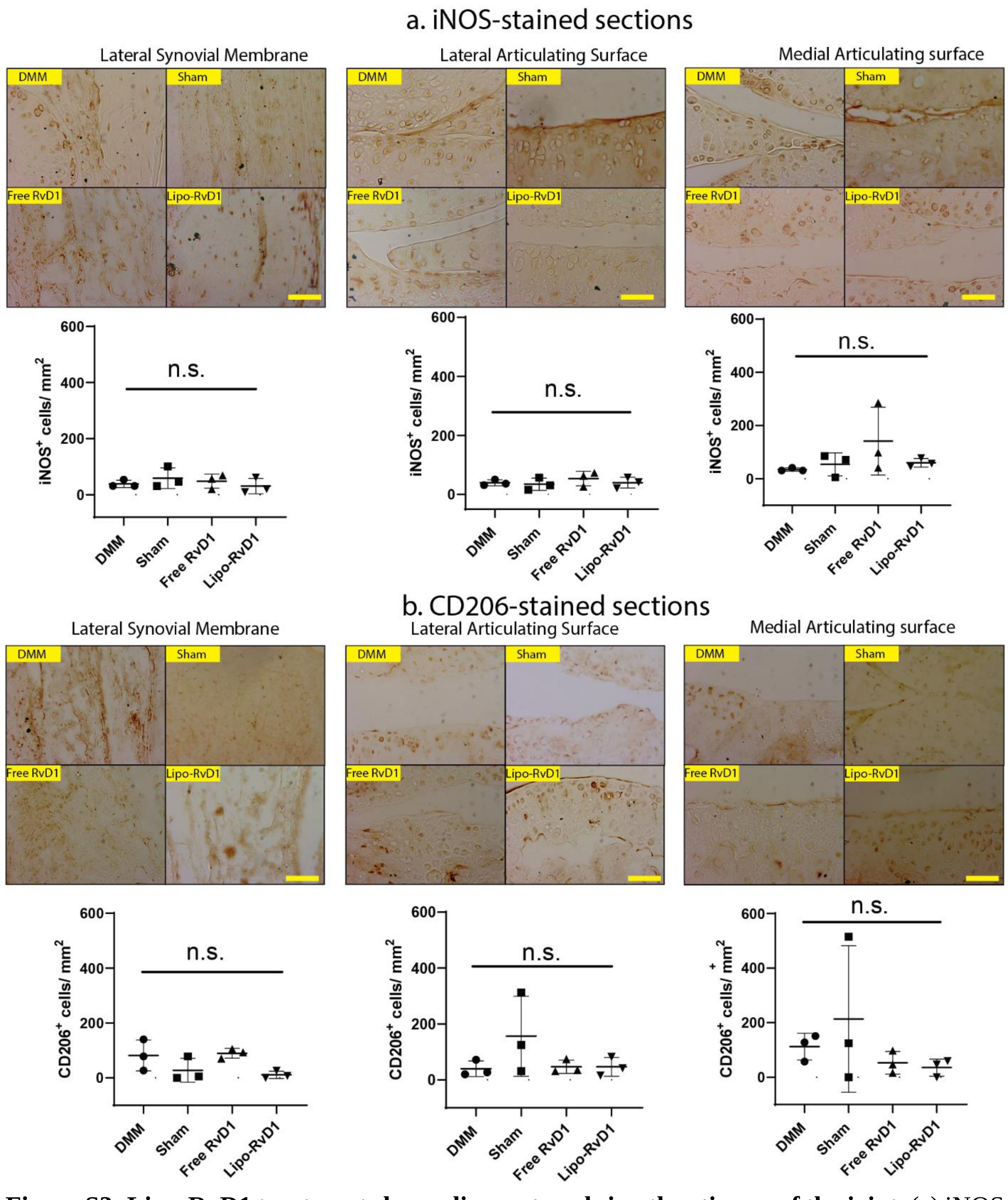


**Figure S3: Lipo-RvD1 treatment shows diverse trends in other tissues of the joint.** (a) iNOS and (b) CD206 stained sections and signal quantification of the respective parts of the joint; n = 3 joints per group. Scale bar 50 µm.

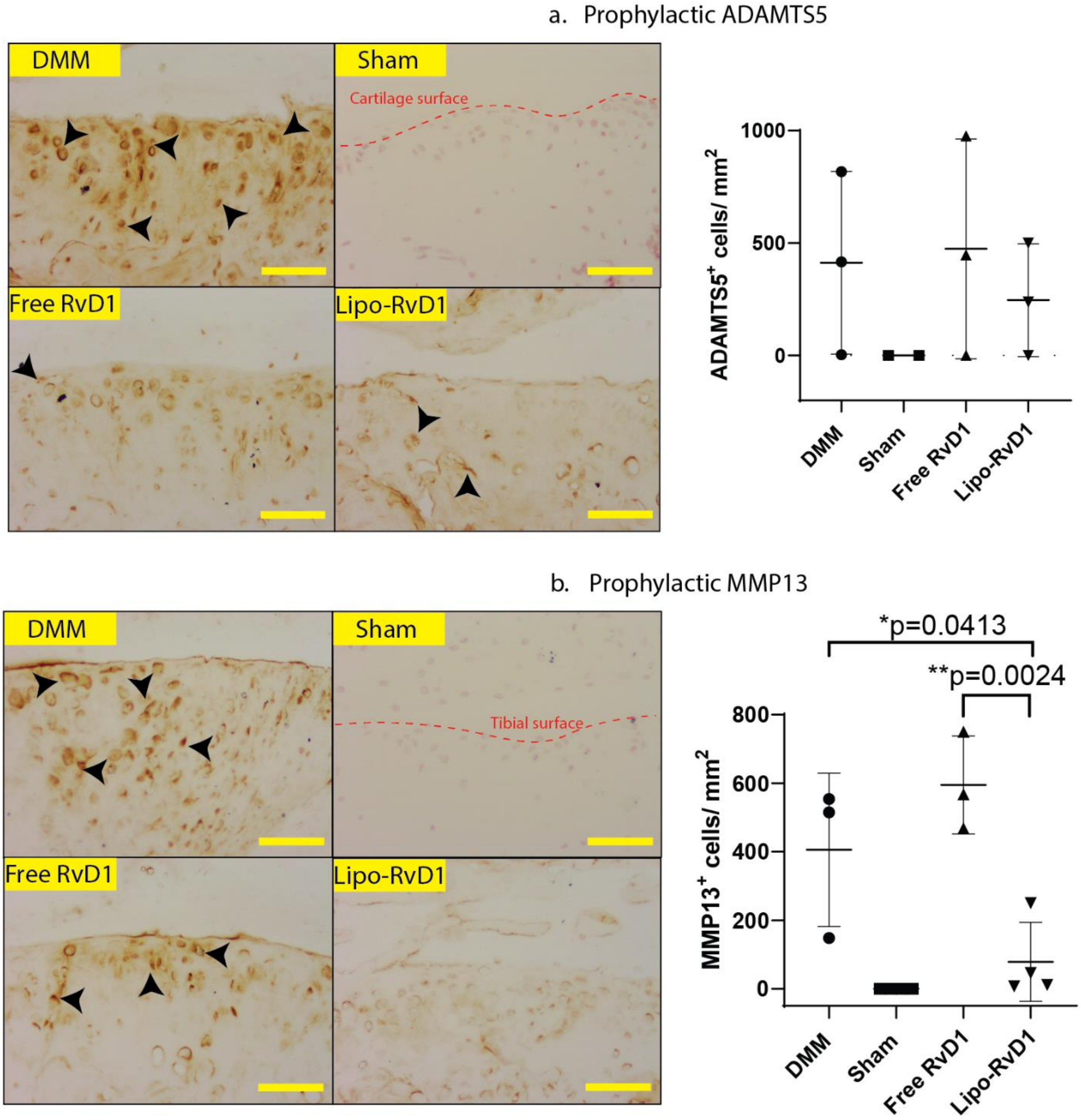


**Figure S4: Lipo-RvD1 when administered prophylactically reduces the expression of catabolic mediator enzymes**. Immunohistochemical images of sections stained for expression of the catabolic markers and its quantification, (**a**) ADAMTS5 and (**b**) MMP13. Scale bar 50 µm.

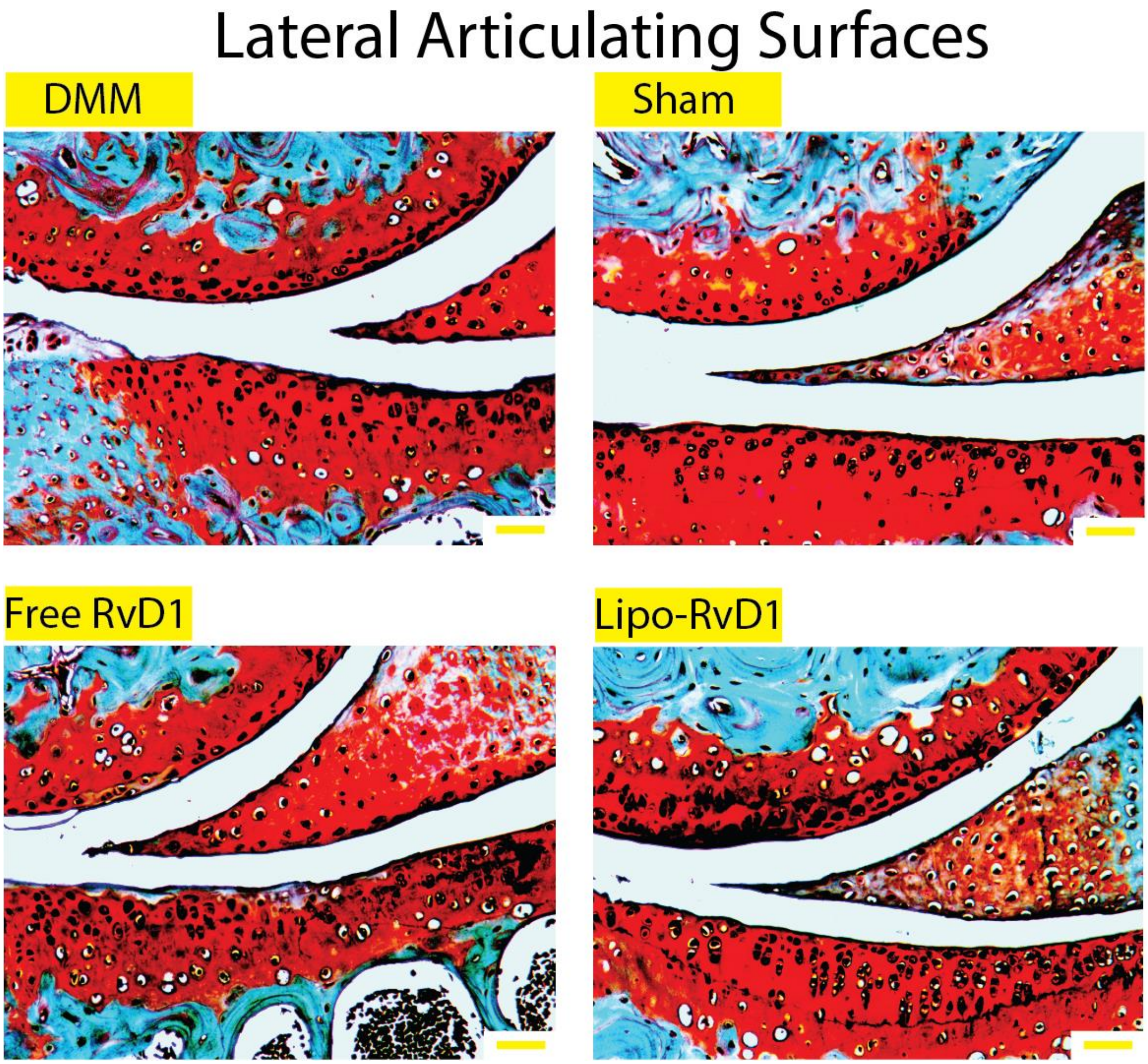


**Figure S5: Cartilage damage does not extend to lateral regions of the affected joints.** Lateral condyles of safranin O-stained joint sections of the respective groups in the therapeutic treatment regimen. Scale bar 50 µm.

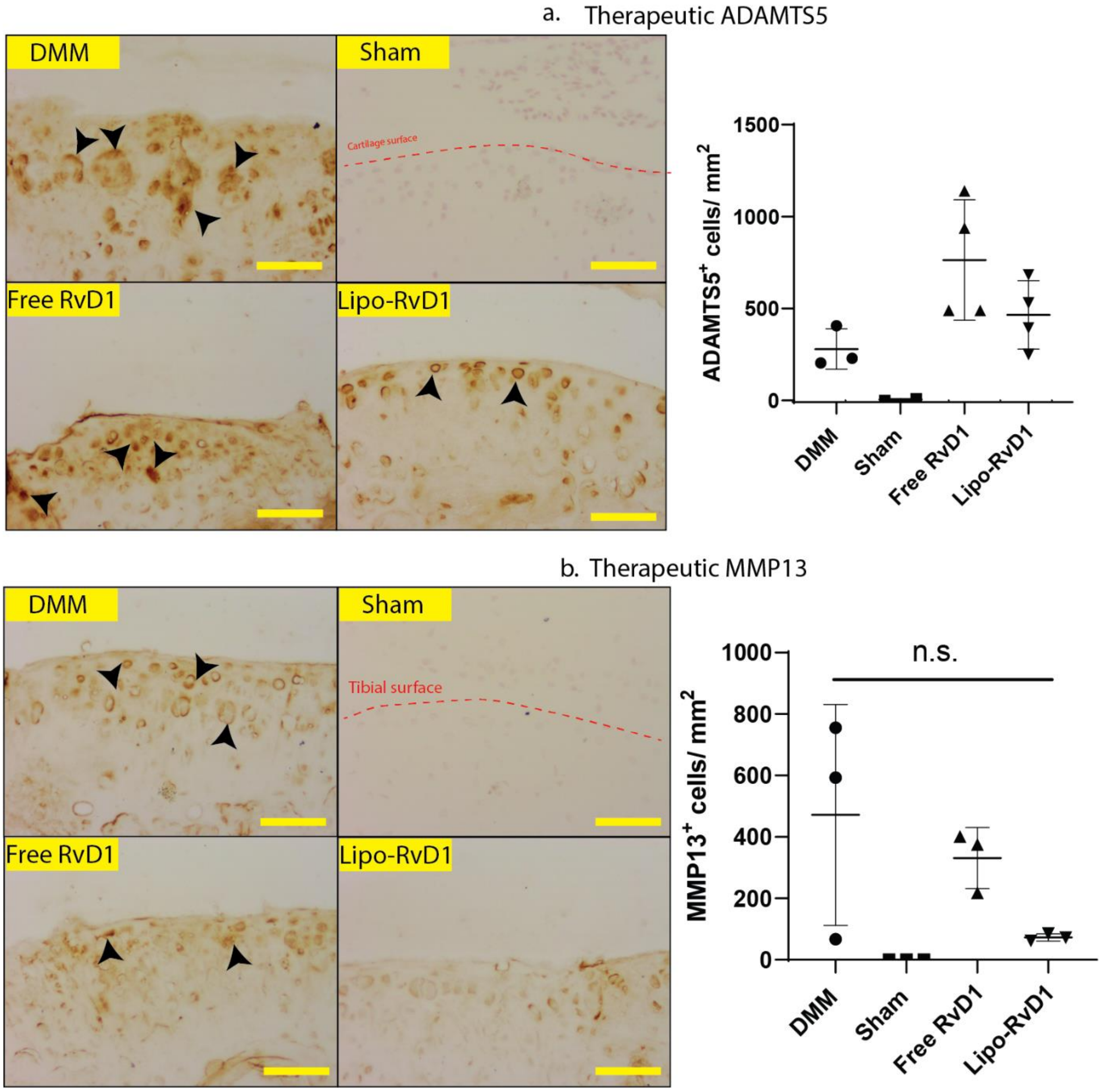


**Figure S6: Lipo-RvD1 when administered therapeutically reduces the expression of catabolic mediator enzymes**. Immunohistochemical images and quantification of sections stained for expression of the catabolic markers and its quantification, (**a**) ADAMTS5 and (**b**) MMP13; n=3-4 joints per group. Scale bar 50 µm.

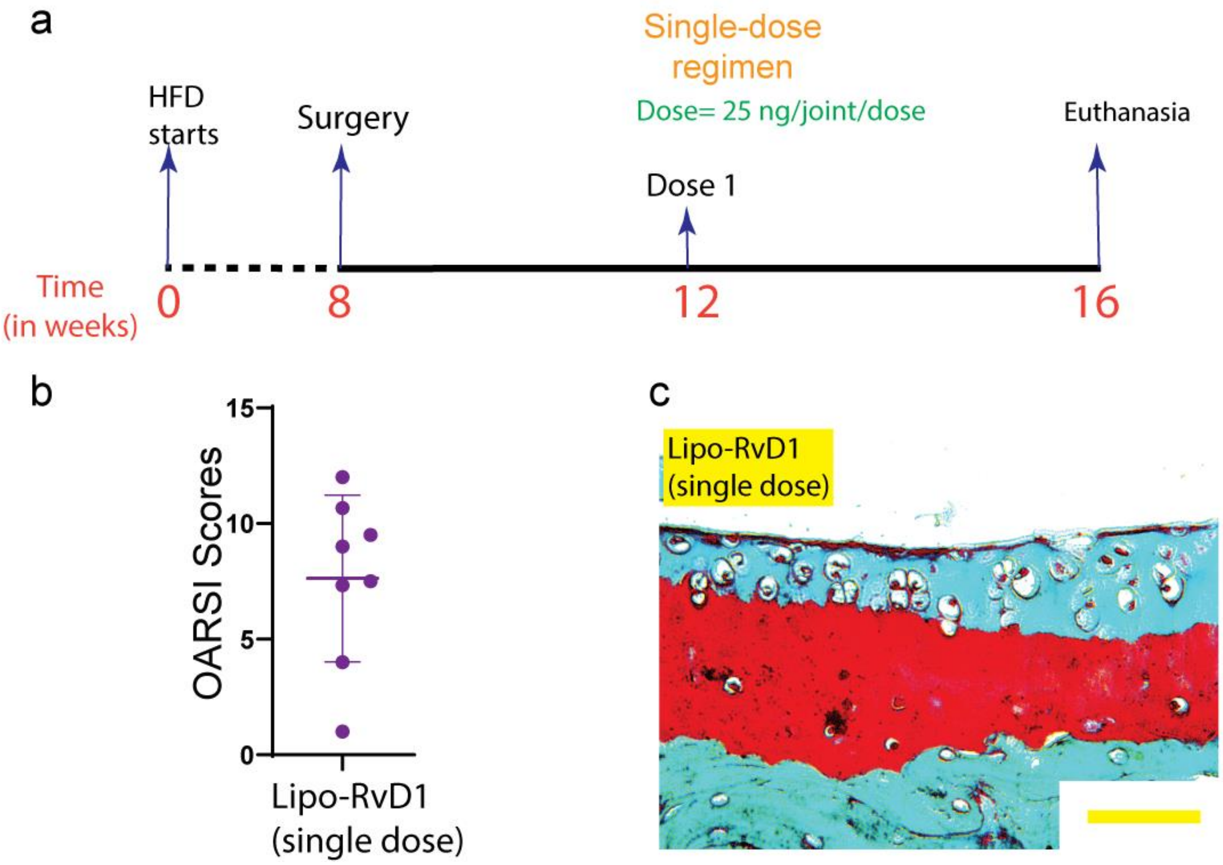


**Figure S7: Single dose of lipo-RvD1 is not sufficient to protect against joint damage.** (**a**) Timeline of the study. (**b**) OARSI scores of histological sections depicting overall joint health; n=8 joints. (**c**) Safranin-O-stained histological sections. Scale bar 50 µm.

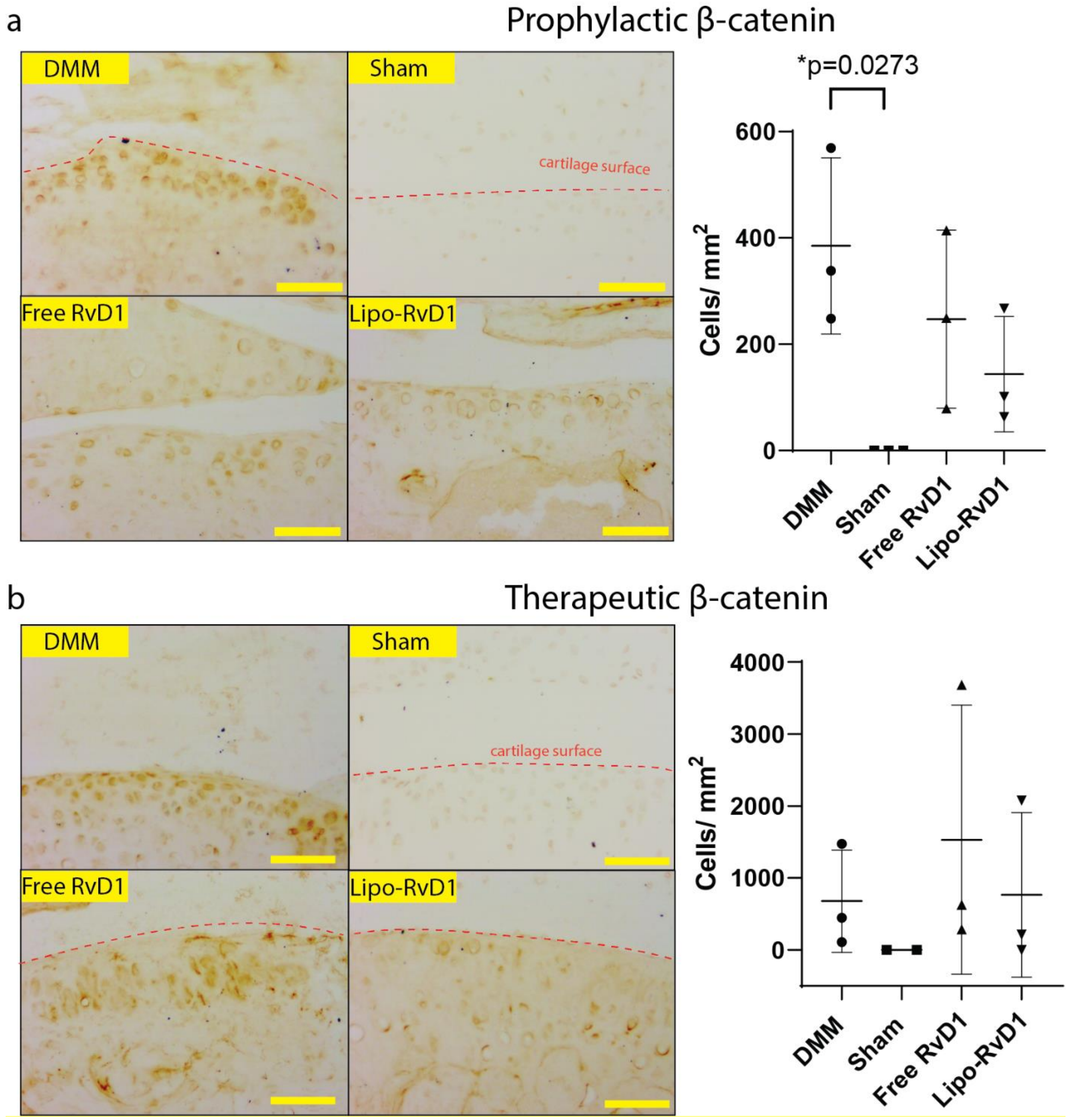


**Figure S8: Lipo-RvD1 administration reduces the β-catenin signaling in joints**. Immunohistochemical images of sections stained for expression of β-catenin and its quantification in (**a**) prophylactic and (**b**) therapeutic regimens. For **a**, *$p$<0.05 between the respective groups indicated in the figures using ANOVA followed by Tukey's posthoc test. Values are expressed as mean ± SD. Scale bar 50 µm.

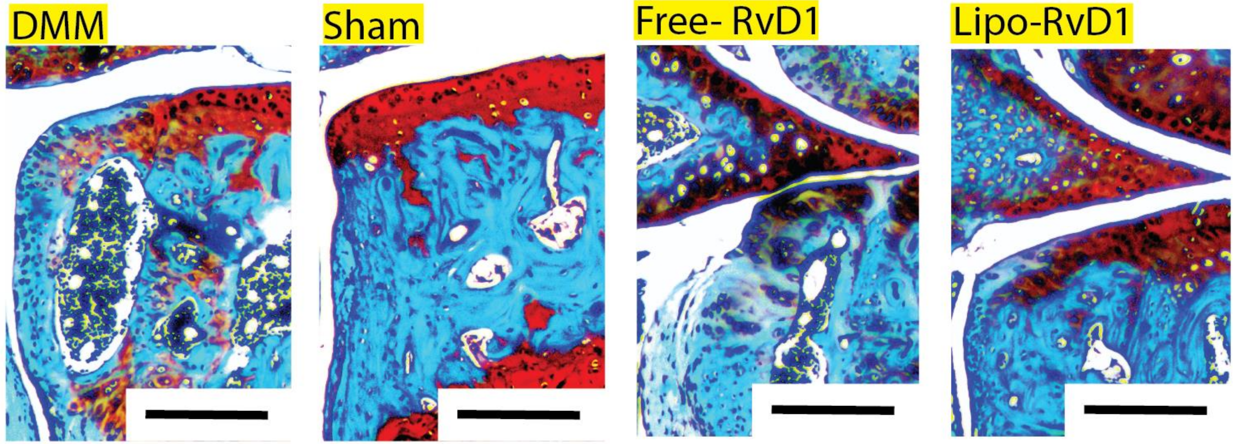


**Figure S9: High-fat fed mice joints do not show development of chondrophytes.** Tibial plateaus of joints administered respective treatments. Scale bar 200 µm.
